# The importance of population growth and regulation in human life history evolution

**DOI:** 10.1101/001404

**Authors:** Ryan Baldini

## Abstract

Explaining the evolution of human life history characteristics remains an outstanding problem to evolutionary anthropologists. Progress is hindered by common misunderstandings of how selection works in age-structured populations. I review two important results of life history theory related to demography. First, different life history strategies evolve under density-independent and density-dependent population growth. Second, and more poorly appreciated, different kinds of density-dependence also select for different life history strategies; assuming zero population growth alone is insufficient to determine the optimal strategy. I show that these facts are more than methodological niceties by reanalyzing the model by Kaplan et al. (2000) and showing that the results depend strongly on the form of population regulation assumed. This analysis suggests that progress in human life history theory requires better understanding of the demography of our ancestors. I close with a discussion of empirical implications.

## 1 Introduction

The unusual life history characteristics of humans provide a unique challenge to evolutionary anthropologists. Humans have a distinctive life history even among our closest living relatives: compared to other primates, we endure a long juvenile period of intensive growth and learning as well as extreme dependency; we mature and reproduce late; we have long reproductive careers with short interbirth intervals; and we even enjoy a long post-reproductive lifespan (Kaplan et al., 2000; Mace, 2000). This pronounced life course has attracted the attention of many theorists, and consequently some novel, human-specific theories have been proposed (Hawkes et al., 1998; Kaplan et al., 2000; Kaplan and Robson, 2002) along with more traditional applications of life history results studied in non-humans (Hill, 1993; Mace, 2000; Kaplan and Gangestad, 2005).

Even sophisticated models of life history evolution, however, sometimes make methodological errors, or rest on specific (and sometimes unrecognized) assumptions that greatly affect the results. In this paper, I discuss one important, under-appreciated aspect of life history evolution: the effect of population growth and regulation. I review theory showing that selection under different forms of population regulation can favor very different kinds of reproductive strategies. First, I review the well-known result that selection in exponentially growing populations often favors different strategies than in stationary populations. Second, I review the less well-known theory showing that even different kinds of density dependence, all of which lead to zero population growth, can favor different life history strategies. This implies that it is not in general correct to assume zero population growth and then maximize a density-independent function of lifetime reproductive success, as theorists sometimes do (e.g. Roff (1992); Stearns (1992)). Rather, one must specify precisely how vital rates depend on population density. Different functions usually produce different outcomes of selection.

To demonstrate the importance of these effects, I reanalyze the model in an oft-cited paper on the evolution of human life histories: Kaplan et al.’s (2000) “A theory of human life history: diet, intelligence, and longevity.” I show that Kaplan et al.’s original model implicitly assumed that density dependence acts only on fertility, independent of age (implying that mortality, for example, does not depend on population density). Their conclusions depend strongly on this assumption, as simple alterations to the form of density dependence produce different outcomes, even while stated assumptions remain intact. This sensitivity to the precise form of population regulation implies that we need to better understand the demographic history and prehistory of our species if our models are to provide specific, realistic predictions.

The next section reviews the theory of maximization in life history evolution. I show what is maximized under various forms of population regulation, which are later applied to the model in Kaplan et al. I devote particular attention to the fact that selection does not, in general, maximize the lifetime number of offspring produced by a strategy. Section 3 describes the Kalpan et al. model, and then applies the theory of section 2. Section 4 closes with a discussion of how empiricists can make use of the theory discussed here.

## 2 What does selection maximize in age-structured populations?

Models of life history evolution often assume that natural selection maximizes some quantity. The goal of this section is to clarify which quantities (if any) are maximized under various situations. I devote extra attention to showing why *R*_0_, the expected lifetime number of offspring, is not maximized under density independence, and is insufficient to determine the action of selection even under density dependence.

### What is meant by maximization?

I define here precisely what I mean when I say that selection maximizes some quantity. The definition is equivalent to that implied by many models in behavioral ecology (including Kaplan et al. (2000)).

Consider a collection of heritable life history strategies that faithfully reproduce over time. For concreteness, it is useful to imagine that these strategies are coded by separate alleles in a haploid system of inheritance (as in Grafen’s “phenotypic gambit” (1984)). Suppose that, to each strategy, a single number measuring some quantity can be assigned. Let *w* denote the value of this quantity; *w_i_* is the quantity for strategy *i*. If the strategy *a* with *w_a_* can invade, replace, and exclude another strategy *b* if and only if *w_a_ > w_b_*, then we say that “selection maximizes *w*.” Then, as long as all possible strategies eventually arise (via mutation or migration), the strategy with the highest *w* will eventually dominate the population to the exclusion of all others. I’ll call *w “*fitness,” but note that fitness is also used to refer to the instantaneous growth rate of strategies, as well properties of individuals (e.g. long-term genetic contribution to a population).

A crucial point to keep in mind throughout the paper is that selection does not maximize any quantity, in general. Maximization can be precluded by some forms of density-dependent and stochastic population growth (Turelli and Petry, 1980), details of the genetic architecture (even sexual reproduction alone can preclude maximization in some cases), and especially frequency-dependent selection (i.e. when the fitness of strategies depends on the frequencies of other strategies in the population; Rice, 2004). All of these undoubtedly affect human evolution, so the models analyzed here are surely incomplete. For example, if much of the information learned by human juveniles is culturally transmitted, then models that consider the evolution of delayed reproduction due to intensive juvenile learning should ideally include co-evolution of social learning strategies with life history. Social learning strategies always have frequency-dependent fitness, so the maximization methods treated here would then be irrelevant; ESS models would be necessary. Still, I ignore these details in this paper, as the goal is to revisit the phenotype-focused maximization methods of life history evolution, which rarely consider these complications. Thus, from now on, I consider only asexual inheritance of strategies, deterministic population growth, and no frequency dependence. For this simple case, it is sometimes possible to derive maximization principles; I review these just below. Readers can find detailed derivations of most results in Charlesworth (1994).

### Maximization under density-independent population growth

Under density-independent growth in asexual populations, and without environmental stochasticity, selection maximizes *r*, the strategy’s exponential rate of increase. *r* is a function of the fertility and mortality rates. It is found by solving the Euler-Lotka equation

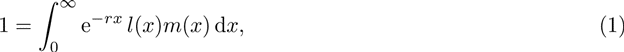

where *l*(*x*) gives the probability of surviving to age *x*, and *m*(*x*) the fertility rate at age *x*. Strictly speaking, populations only grow at rate *r* when the stable age distribution (or SAD) is reached, but convergence to the SAD virtually always results under human vital rates (Caswell, 2001).

This maximization rule is simple to understand. With asexual reproduction and density independence, strategies can be imagined as entirely separate populations, each growing at a different exponential rate. The one that grows fastest eventually reaches frequency 1, relative to all others.

### *R*_0_ is not maximized under density independence

Selection does not maximize the expected lifetime number of offspring (*R*_0_) under density-independent growth. For most, this fact is not immediately obvious, so I provide a simple demonstration. Imagine two asexual life history strategies, *A* and *B*. All individuals survive for exactly two seasons and produce two offspring, but *A* and *B* differ in their timing of reproduction. Individuals of strategy *A* produce one offspring in both their first and second seasons, while individuals of strategy *B* produce both offspring in their second season only.

Table 1 shows the result of selection, beginning with a population composed of one newborn individual of each strategy. Strategy *A* grows faster than *B* and eventually approaches frequency 1 in the population. The reason is that *A* reproduces much earlier than *B*, and thus has a shorter generation length and ultimately a higher *r*.

**Table 1:**
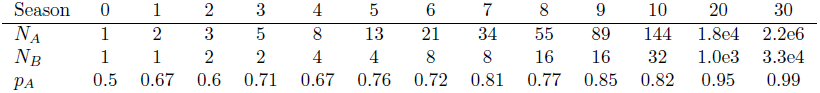
*N_A_* and *N_B_* give the population sizes for strategies *A* and *B*, respectively. *p_A_* is the frequency of strategy *A* among the total population. The frequency of *p* fluctuates as it approaches 1 because strategy *B* does not converge a unique age distribution and growth rate.

| Season | 0 | 1 | 2 | 3 | 4 | 5 | 6 | 7 | 8 | 9 | 10 | 20 | 30 |
| --- | --- | --- | --- | --- | --- | --- | --- | --- | --- | --- | --- | --- | --- |
| $N_A$ | 1 | 2 | 3 | 5 | 8 | 13 | 21 | 34 | 55 | 89 | 144 | 1.8e4 | 2.2e6 |
| $N_B$ | 1 | 1 | 2 | 2 | 4 | 4 | 8 | 8 | 16 | 16 | 32 | 1.0e3 | 3.3e4 |
| $p_A$ | 0.5 | 0.67 | 0.6 | 0.71 | 0.67 | 0.76 | 0.72 | 0.81 | 0.77 | 0.85 | 0.82 | 0.95 | 0.99 |

A useful approximation of *r*, when *r* is small, is given by the quantity

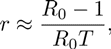

where *T* measures the average age at reproduction among individuals of the strategy (Charlesworth, 1994).^1^ All other things being equal, selection favors smaller *T* in growing populations. In our example, *T_A_* = 1.5, whereas *T_B_* = 2, which explains why *A* quickly outgrows *B*. Although the example provided here is contrived, recent findings suggest that age at reproduction may well be an important factor in human evolution. See section 4 for a discussion of some empirical examples.

### Maximization under density-dependent population growth

Density-dependent population growth, under which vital rates depend on population density, greatly complicates selection dynamics. Density dependence can be extremely complicated, and as a result, selection generally does not maximize any quantity: multiple strategies may mutually exclude each other, or may coexist in a polymorphic population. Charlesworth (1994) derived one reasonable case under which maximization does occur, and I will treat this case throughout. Suppose that all strategies are sensitive to the same function of the number of individuals in some age group. Let us simply call this quantity *N*, the critical population size. *N* could be the total population size, the total number of juveniles alone, or any increasing function of the population size across ages (e.g. *N* could be a weighted sum of the different age classes, with older age classes carrying more weight). Assume furthermore that *l* and *m* are non-increasing functions of *N*; population density only adversely affects survival and fertility (at least in the region of the equilibrium). Then selection maximizes *N̂*, the equilibrium *N* found by solving the equation

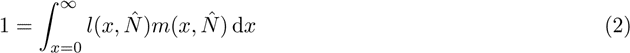

*N̂* can be considered the carrying capacity of the strategy with respect to the critical population size. This is still only true if all strategies approach a stable *N̂* over time; population cycles and chaos preclude optimization.

This result can be understood as follows. Suppose a resident strategy has achieved equilibrium such that *N*, the critical population size, has approach *N̂*_1_. Now suppose a new strategy arises in the population, which is characterized by a slightly larger *N̂*_2_ - that is, if it were to grow by itself in a separate environment, it would achieve higher *N* at equlibrium than the resident strategy. Since *l* and *m* are non-increasing functions of *N*, and *N̂*_1_ < *N̂*_2_, it follows that the new strategy grows upon introduction. The resulting total population growth implies negative growth for the resident strategy. Eventually the new strategy replaces the old entirely, and when the dust clears, *N̂* has increased.

### *R*_0_ is insufficient to determine the outcome of selection under density dependence

The misuse of *R*_0_ for populations held in check by density-dependent growth is more subtle than the problem discussed for density independence. The Euler-Lotka equation (1) shows that whenever a resident strategy has *r* = 0, so that the population is stationary, then *R*_0_ = 1. A new strategy invades this population if its growth rate is positive upon introduction. One can show that this is equivalent to the condition that *R*_0_ *>* 1 for the invading strategy, or that the expected lifetime offspring production of the new strategy is above replacement upon introduction. *R*_0_ can then be considered a measure of fitness in the sense that it completely determines whether a new strategy can invade (Hastings, 1978), as well as determining the instantaneous growth rate of a novel strategy (Charlesworth, 1994). That is, selection maximizes *R*_0_ in the following, local sense: when the *optimum* strategy is at carrying capacity, *R*_0_ = 1 (by definition), and any *nonoptimal* strategy would have *R*_0_ *<* 1 upon introduction. *R*_0_ is not maximized in the global sense as used in this paper (and as implied in Kaplan et al. (2000) and other maximization analyses): strategies do not have a fixed *R*_0_, so global maximization as defined before is impossible.

The error of using *R*_0_ arises when theorists then construct fertility and mortality functions *that are entirely independent of density*, compute a unique *R*_0_ from these, and globally maximize *R*_0_ with respect to some life history variable. That is, given some strategy *v* and demographic functions independent of *N*, they seek the *v̂* that solves the equation

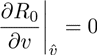

(They also check that this is a maximum instead of a minimum.) By not specifying how *l* and *m* depend on population density, however, this method invariably violates the constraint that *R*_0_ = 1 at equilibrium. The correct way to work with *R*_0_ directly is to specify how *R*_0_ depends on *N*, and then solve the *two* equations

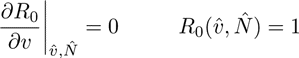

This is equivalent to maximizing *N̂*, as the above theory demands. Specifying the form of population regulation is crucial, because it strongly affects the outcome of selection. We will see this in the reanalysis of Kaplan et al. (2000) below.

Examples of the erroneous method applied to nonhuman evolution can be found in both Stearns’ (1992) and Roff’s (1992) book-length reviews of life history evolution. Figure 6.23 in Stearns (concerning the optimal age at maturity, pp. 145-147) clearly reveals the error of this method: in the example given, the optimal strategy has *R*_0_ *>* 20, which is clearly inconsistent with the requirement that *R*_0_ = 1 under zero population growth! Neither Stearns nor Roff writes that the methods they use only happen to be valid if density dependence acts on fertility or newborn mortality alone (Charlesworth, 1994; I review this below). Kaplan et al. make the same implicit assumption.

## 3 The coevolution of age at maturity and investment in survival

I now shift attention to the model by Kaplan et al. (2000). I review the assumptions and claims of the model, and then reanalyze it assuming various forms of population regulation.

### The model

The model in Kaplan et al. (2000, p. 165) treats the coevolution of two life history characteristics: age at maturity and investment in survival. Individuals experience two life stages: a juvenile, pre-reproductive stage, and an adult, reproductive stage, which begins at age *t*. In both stages, individuals invest some proportion *λ* of available energy in mortality reduction. Among juveniles, the remaining energy is devoted to the development of embodied capital (growth and learning). Among adults, the remaining energy is devoted to reproduction. Only *t* and *λ* evolve in this model.

The instantaneous death rate, *µ*, is constant across ages for any given strategy. Since *λ* measures investment in mortality reduction, *µ* is a decreasing function of *λ*: strategies with high *λ* live longer. To allow for external effects on mortality, the parameter *θ* is introduced to quantify the extrinsic mortality risk. *µ* is an increasing function of *θ*.

Kaplan et al. do not provide fertility functions, but they do provide a term that represents the energy invested in fertility. I simply change this to represent fertility outright, so that an *m* function is recovered. Fertility at age *t* is equal to the embodied capital produced up to that point, multiplied by 1 − *λ* (as the rest of the energy continues to be invested in survival). Let the embodied capital (and therefore fertility) at age *t* be denoted by *P* (*t, λ*, ∈). *P* is an increasing function of *t*: more time in maturity allows for more embodied capital. It is a decreasing function of *λ*, as this energy is lost to survival investment. *∈* is an ecological parameter that measures the ease of the environment with respect to fertility: all other things being equal, greater *∈* implies higher fertility. Finally, Kaplan et al. assume that energy production grows at an exponential rate *g* after maturity due to skills and knowledge acquired from experience during the reproductive stage. I translate this energy production directly to fertility.

### The claimed results

Kaplan et al. claim six general results with regard to evolution in *t* and *λ*. Let *t̂* and λ̂ be the optimal values of age at maturity and investment in mortality reduction, respectively. Then Kaplan et al. claim

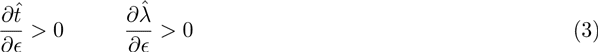

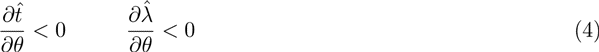

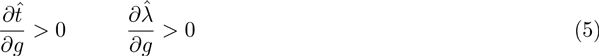

Inequalities (3) say that, all other things being equal, niches more hospitable to reproduction favor later maturity and greater investment in mortality reduction. Inequalities (4) say that as extrinsic mortality increases, selection favors earlier maturity and lesser investment in mortality reduction. Inequalities (5) say that a greater growth rate of energy production (and therefore fertility) after maturity selects for later maturity and greater investment in mortality reduction.

Kaplan et al. emphasize not only the directional effects of the ecological parameters, but also the positive coevolutionary relationship between age at maturity and investment in survival. For parameter change, they found that the age at maturity and the investment in survival always increased or decreased together (i.e. for each line, the derivatives have the same sign).

### Reanalysis

#### Forms of population regulation

The main problem with the original analysis in Kaplan et al. is a failure to specify the form of population regulation, which strongly effects the outcome of selection. Specifically, they make the error described in section two: they claim zero population growth without specifying precisely how this occurs. Here I describe specific forms of population regulation applied to the model. I generalize by allowing one case of nonzero population growth, as well as various forms of density dependence.

Consider four forms of population regulation:

1. Density-independent (exponential) growth
2. Density-dependent fertility
3. Density-dependent mortality
4. Density-dependent fertility *and* mortality

In case 1, all fertility and mortality rates are fixed quantities, independent of population density. Selection maximizes *r* here, whereas it maximizes *N̂* for all other cases.

In case 2, fertility across all ages depends on population size via the same multiplier. In particular, I assume that 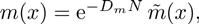 where 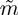 is the fertility under zero population density and *D_m_* quantifies the extent to which fertility is adversely affected by population density. Thus, *m* → 0 as *N* → ∞. Survival rates are independent of density.

In case 3, the same form of density dependence applies to mortality, rather than fertility. That is, *l*(*x*) = 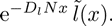 Fertility is independent of density.

In case 4, both fertility and mortality are density dependent, using the forms of cases 2 and 3.

In all density-independent cases, the *D* parameters are assumed to be fixed. That is, all strategies are affected by density equally. I assume this only for simplicity, as different strategies could realistically vary in their susceptibility to crowding (this is the foundation of *r* vs. *K* selection theory). Fortunately, this assumption also leads to several useful simplifications (Charlesworth, 1994). First, one can show that cases 1 and 3 favor the exact same strategy for *l* and *m*. This follows because *r* and *D_l_N̂*, which are respectively maximized in cases 1 and 3, take the same functional form in the Euler-Lotka equation. Second, maximizing *N̂* in case 2 is equivalent to maximizing the strategy with the largest *R*_0_ under zero population density. This implies that Kaplan et al.’s analysis (and others) implicitly and unknowingly assumed that density-dependence acts only on fertility, and equally across all ages. Finally, the results of case 4 lie somewhere between the results of cases 2 and 3. As intuition suggests, if *D_m_ ≫ D_l_*, such that density dependence acts more strongly on fertility than mortality, then the results of case 4 closely resemble that of case 2, and vice versa.

These simplifications imply that we need only consider two cases: cases 1 and 3 together (as they favor the same strategy), and case 2. The results of case 4 lie somewhere between these two. These simple differences in population regulation lead to very different optimal life history strategies.

#### Cases 1 and 3

Under density-independent population growth, we have the following demographic equations:

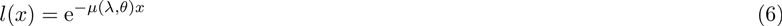

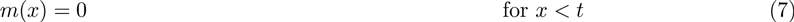

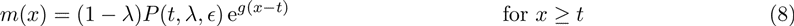

Under density-dependent mortality, *l*(*x*) is multiplied by the term 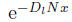 as described above.

Under density-independence, a strategy’s growth rate is found by plugging the functions (6), (7), and (8) into equation (1). Doing this and simplifying produces^2^

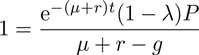

Solving for *r* produces

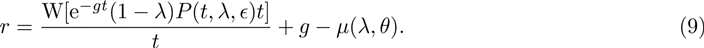

W is the product-log function: W[*z*] gives the solution for *w* in *z* = *e^w^*. Though W is a complicated function, it is useful to know that *W* [*z*] *>* 0 for *z >* 0 and d*W/*d*z >* 0, i.e. the product-log is an increasing function of its argument.

An invading strategy will only invade if the *r* of the invading strategy is greater than that of the resident strategy. If mutant strategies both *t* and *λ* tend to differ from their resident values by a small amount, then the direction of evolution is predicted by the derivatives of *r* with respect to both strategy variables. Furthermore, any internal equilibrium in *t* and *λ* must satisfy the condition that both derivatives equal 0. Thus we search for optima via the simultaneous solution of d*r/*d*t* = 0 and d*r/*d*λ* = 0. Simplifying these gives us

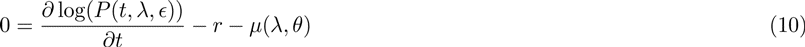

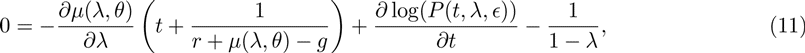

where *r* is given by equation (9). (We must also check that the solution actually maximizes, rather than minimizes, *r*.) Note how these differ from equations (2) and (3) in Kaplan et al. (2000) (reproduced below in (14) and (15)).

The analogous equations for the case of density-dependent mortality are found by replacing *r* in the above equations with *D_l_N̂*, subject to the constraint that *N̂* implies 0 population growth as in equation (2). These two models favor the same strategy at equilibrium, so all further results apply to both.

Due to the complexity of these equations, I have been unable to derive any general results about parameter effects, leaving claims (3), (4), and (5) difficult to evaluate in general. To proceed I assume explicit functions for *P* and *µ* and seek numerical solutions. I assume the functions

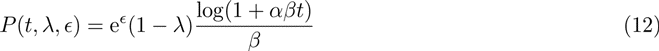

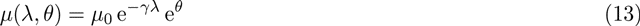

Equation (12) implies that fertility at maturity is diminishing-returns function of age at maturity. The parameter *α* measures the initial rate at which embodied capital is acquired during the juvenile period. *β* is the diminishing-returns parameter: with higher *β*, the more quickly returns to learning and growth diminish. In equation (13), *γ* determines the rate at which investments in survival (*λ*) decrease mortality risk. The remaining parameters were defined above and satisfy the assumptions of Kaplan et al. (2000).

Equations (12) and (13) can be substituted into equations (10) and (11), which can then be solved numerically. While this procedure cannot prove general results, we readily find that most of Kaplan et al.’s claims do not hold in this case (see figure 1 for examples). Under all parameter values numerically investigated (see appendix A), I find that, contrary to inequalities (3), *t̂* and λ̂ both *decrease* as *∈* increases; niches more hospitable to reproduction prompt earlier maturity and lesser investment in survival. Contrary to inequalities (4), *t̂* and λ̂ both *increase* with *θ*; niches with greater extrinsic mortality risk favor later maturity and greater investment in survival. Finally, while λ̂ does increase with *g* as Kaplan et al. predicted, *t̂* decreases; niches in which production continues to grow in adulthood favor earlier maturity. This latter finding also shows that *λ* and *t* may not evolve in the same direction.

**Figure 1:**
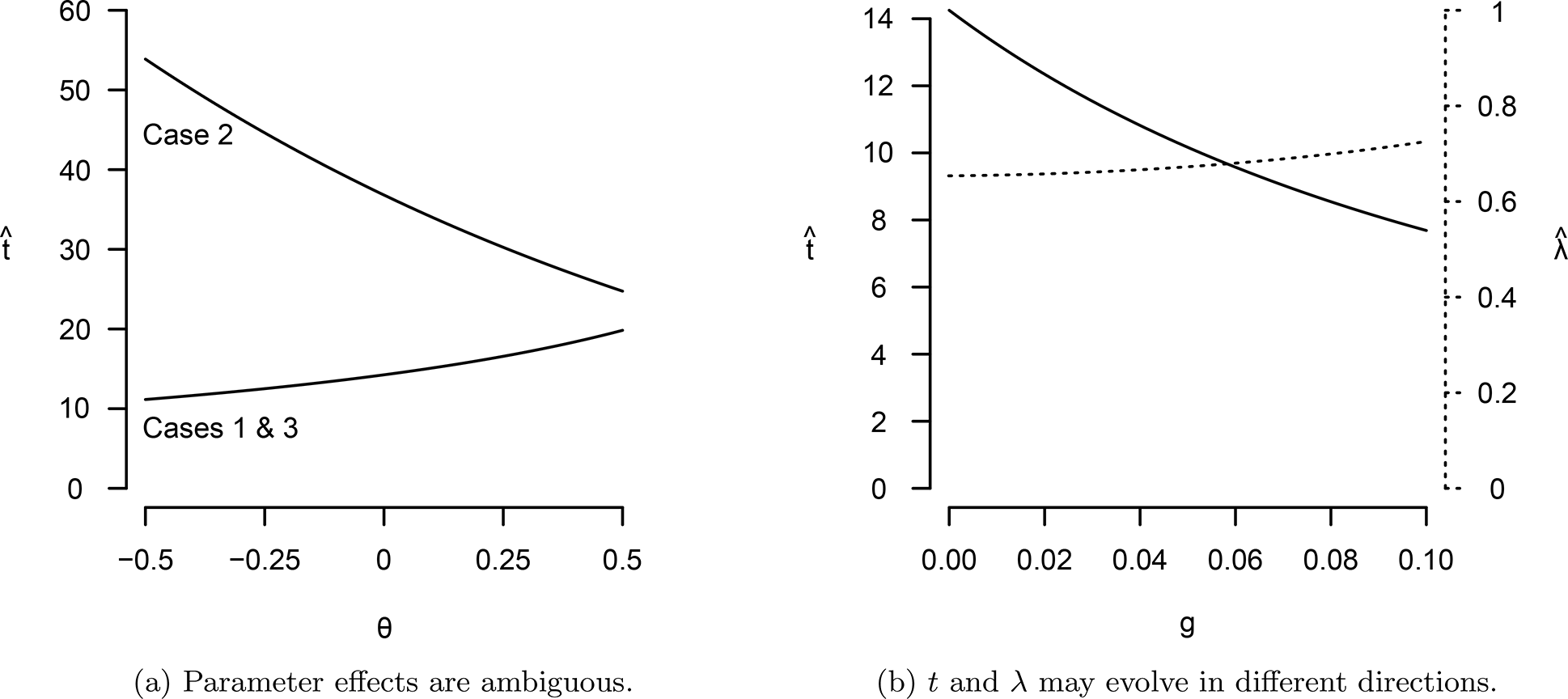
Contradictions of Kaplan et al.’s (2000) claims. (a) The optimal age at maturity, *t̂* may either decrease or increase with extrinsic mortality risk, *θ*, depending on the form of population regulation. (b) Under cases 1 and 3, *t* and *λ* evolve in different directions when *g* increases. The solid line depicts *t̂* while the dashed line depicts λ̂ For both (a) and (b), *α* = 0.1, *β* = 0.2, *µ*_0_ = 1, *γ* = 5, *∈* = 0, *θ* = 0, *g* = 0, except when the latter two vary.

#### Case 2

Repeating the same process as above, but assuming that population density only affects fertility, we arrive at the equilibrium conditions

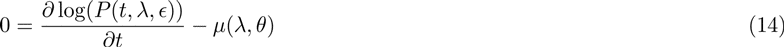

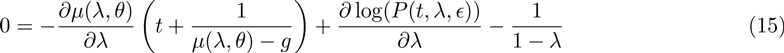

These are equivalent to equations (2) and (3) in Kaplan et al. (2000). This is expected because this form of density dependence maximizes *R*_0_ under zero population density.

Curiously, some of Kaplan et al.’s results do not hold even in this case, suggesting that the authors made assumptions that were not printed in the paper. Again assuming functions (12) and (13), I first find that *∈* drops out entirely; the extrinsic costs of fertility have no effect on the equilibria. Second, numerical calculations show that *λ* evolves upward with *θ*, which is opposite to *t*. Finally, both *t* and *λ* evolve upward with *g*, as Kaplan et al. predict.

### What can we conclude?

All told, my analysis casts doubt on the general claims of Kaplan et al. (2000). Even the simple cases considered above (recall that density affected all ages equally, and *D*’s did not vary between strategies) showed great diversity in the outcome of selection. More complicated forms of population regulation would probably produce still more widely varying outcomes. Thus any strong claims like inequalities (3), (4), and (5) seem unlikely to hold in general; there may be *some* form of density-dependence for which *any* conclusion does not hold.

Still, to avoid leaving the reader with the unsatisfying message that “anything can happen,” I conducted a systematic search for general parameter effects (appendix A describes the methods). I found two consistent effects: first, increasing *α*, the initial rate at which embodied capital grows, always favors earlier maturity and lesser investment in survival. Second, increasing *g*, the rate at which production (and, hence, fertility) grows after maturity, always favors greater investment in survival. Fortunately, these conclusions seem compatible with Kaplan et al.’s general argument. If the transition to the intensive human foraging niche caused slower initial rates of growth and learning (due to the initial difficulty of acquiring necessary skills), and allowed for greater production growth during adulthood, then this may well have contributed to the evolution of the human extended life history. Other ecological effects, unfortunately, remain ambiguous. For example, if the niche transition also implied higher extrinsic mortality due to the difficulty of acquiring resources, then this could have selected for either shorter or longer life histories, depending on the form of population regulation (figure 1(a)).

The lack of generalizable results implies that we need to shift our focus away from abstract evolutionary models and toward understanding the demographic conditions faced by humans throughout the time period during which our unique life history evolved. Knowledge of these details would allow theorists to constrain model space to only the most realistic cases, leading to more pointed predictions. Studies of density-dependent effects in contemporary human populations may be a fruitful starting point. Such research is still rare among humans, but multi-species studies among large mammals consistently find that juvenile morality is more sensitive to environmental stress than is adult mortality (Fowler, 1987; Gaillard et al., 1998; but see Owen-Smith et al., 2005 for a case where adult mortality is driven strongly by predation). Adult fertility also appears to be quite sensitive, although this is sometimes confounded with yearling survival, and does not appear to be independent of the mother’s age (Gaillard et al., 2000). That the effects on fertility and mortality depend on age suggest that even the age-independent models treated in this paper cannot be taken too seriously. On the other hand, if it is true that density dependence acts mostly on fertility and infant survival, then case 2 may turn out to be the closest to reality.

## 4 Empirical implications

### Fitness of strategies vs. individuals

Most of the empirical literature of human life history attempts to estimate fitness as a quantity of the individual. The most sophisticated analyses take the effect of population growth into account by using some measure of reproductive value (Voland, 1990; Goodman et al., 2012; Moorad, 2012). This is a reasonable definition of individual fitness, since one can show that a strategy increases in frequency only if the mean reproductive value at birth of individuals with the strategy is greater than zero.

Some conceptual difficulty arises, however, with the concept of individual fitness. Evolutionary models treat the change in frequency of *heritable strategies* through time, and thus it is the success of strategies that matters. The concept of an individual’s fitness is usually not derived in such models. Furthermore, assigning the proper reproductive value to an individual requires knowing her strategy in the first place - but if strategies are already known, then calculating some measure of individual fitness is redundant. Unfortunately, since we are usually ignorant of mechanisms by which reproductive behaviors are developed and transmitted, the set of strategies is unknown in the first place. The problem only gets worse when behaviors are transmitted non-vertically, e.g. through social learning, for then even the maximization principles derived in population genetics become irrelevant.

Still, individual contributions to the gene pool can provide useful information, so it remains important to estimate these quantities correctly. The method used by Voland (1990) is perhaps the most sophisticated way of estimating fitness for historical populations: the fitness of an individual in the past is estimated by its number of current descendants, each weighted by relatedness and reproductive value. Even this method is not perfect, however, because it assigns equal reproductive value to all individuals of the same age and sex, thereby ignoring heritability of reproductive strategies. Ideally, reproductive value of descendants should depend on genotype (and possibly culturally inherited information) as well. If heritability is weak, however, then it should be so small across distant generations that this method provides a decent approximation. Moorad (2012) uses a different approach, which attempts to infer a potentially unique life history course for each person. Unfortunately the method of strategy construction in this paper is somewhat *ad hoc* and probably not ideal (e.g. it assigns a value of one to the probability of surviving some age class to any individual who happened to survive it, and then only assigns population averages of survival rates to age classes after the individual’s death), but the idea of attempting to infer unknown strategy sets from observed data is intriguing.

### The importance of reproductive timing

It is not yet known how well *R*_0_ approximates fitness in human populations. It is possible that, in most populations, growth is slow enough and generation time long enough that *R*_0_ closely approximates reproductive value. Still, researchers should use the “correct” measure whenever possible. Some recent results suggest that variation in reproductive timing may play an important role in human evolution. Goodman et al. (2013) found that age at first reproduction, rather than completed fertility, was a “crucial” factor in explaining why parents of higher socioeconomic status at the turn of the 20th century produced fewer long-term descendants: wealthy parents and their offspring tend to reproduce later. Moorad (2012) shows that using *R*_0_ as a measure of fitness among mothers in 19th century Utah systematically underemphasizes the importance of selection on early reproduction and overemphasizes that of late reproduction (though correlations between *R*_0_ and reproductive value were quite high). The significance of reproductive timing will crystallize as empirical methods in evolutionary demography continue to improve.

## Footnotes

1 Specifically, 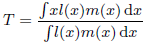

2 For *R*_0_ to remain finite, we require that *g < µ*. I assume this throughout, as do Kaplan et al. (2000)

## A Methods for exploring parameter space

The size of the parameter space precludes exhaustive exploration in this paper. I conducted a restricted search by varying only two parameters at a time, leaving the remaining parameters fixed at baseline values. This allows for identification of first-order interactions between parameters with respect to their effect on *t̂* and *λ̂*. For each interaction (e.g. *α* and *β*), 100 values were uniformly selected from the range of both variables, producing a 100 × 100 matrix and therefore 10,000 separate model equilibria. Equilibria were numerically calculated using the rootSolve package in R.

Baseline values were *α* = 0.1, *β* = 0.2, *µ*_0_ = 0, *γ* = 5, and *∈* = *θ* = *g* = 0. The parameters *α* and *β* were varied by plus and minus an order of magnitude (e.g. 0.01 to 1 for *α*). The remaining parameters involved exponential functions, so were varied over a smaller, additive range. *γ* ranged from 1 to 10; *∈* and *θ* ranged from −5 to 5; and *g* ranged from 0 to 1. Since variation in *µ*_0_ has the same effect as variation in *θ*, *µ*_0_ was not varied.

For model case 1 or 3, I discarded results for any parameter set where *r* or *N̂* was calculated to be less than 0, respectively. The condition was identical across the two cases, so the same results again apply to both. For model case 2, I discarded results for which *N̂* < 0 or *N̂* > ∞. The latter is equivalent to the condition that *g > µ*, which was assumed to not hold in the text, for the same reason. The range of parameter values I used usually surpassed these limits, indicating that the parameter space was well-explored for first-order interactions

## References

Caswell, H., 2001. Matrix Populations Models. Sinauer Associates Sunderland.

Charlesworth, B., 1994. Evolution in Age-Structured Populations. Vol. 2. Cambridge University Press Cambridge.

Fowler, C., 1987. A review of density dependence in populations of large mammals. Current Mammalogy 1, 401–441.

Gaillard, J.-M., Festa-Bianchet, M., Yoccoz, N., Loison, A., Toigo, C., 2000. Temporal variation in fitness components and population dynamics of large herbivores. Annual Review of Ecology and Systematics, 367–393.

Gaillard, J.-M., Festa-Bianchet, M., Yoccoz, N. G., 1998. Population dynamics of large herbivores: variable recruitment with constant adult survival. Trends in Ecology & Evolution 13 (2), 58–63.

Goodman, A., Koupil, I., Lawson, D. W., 2012. Low fertility increases descendant socioeconomic position but reduces long-term fitness in a modern post-industrial society. Proceedings of the Royal Society B: Biological Sciences 279 (1746), 4342–4351.

Grafen, A., 1984. Natural selection, kin selection and group selection. In: Davies, J. R., Krebs, N. B. (Eds.), Behavioural Ecology: an Evolutionary Approach, 2nd Edition. Sinauer Associates Sunderland.

Hastings, A., 1978. Evolutionarily stable strategies and the evolution of life history strategies: I. density dependent models. Journal of theoretical biology 75 (4), 527–536.

Hawkes, K., OConnell, J. F., Jones, N. B., Alvarez, H., Charnov, E. L., 1998. Grandmothering, menopause, and the evolution of human life histories. Proceedings of the National Academy of Sciences 95 (3), 1336– 1339.

Hill, K., 1993. Life history theory and evolutionary anthropology. Evolutionary Anthropology: Issues, News, and Reviews 2 (3), 78–88.

Kaplan, H., Hill, K., Lancaster, J., Hurtado, A. M., 2000. A theory of human life history evolution: diet, intelligence, and longevity. Evolutionary Anthropology: Issues, News, and Reviews 9 (4), 156–185.

Kaplan, H. S., Gangestad, S. W., 2005. Life history theory and evolutionary psychology. In: Buss, D. M. (Ed.), The Handbook of Evolutionary Psychology. Wiley, pp. 68–95.

Kaplan, H. S., Robson, A. J., 2002. The emergence of humans: The coevolution of intelligence and longevity with intergenerational transfers. Proceedings of the National Academy of Sciences 99 (15), 10221–10226.

Mace, R., 2000. Evolutionary ecology of human life history. Animal Behaviour 59 (1), 1–10.

Moorad, J. A., 2012. A demographic transition altered the strength of selection for fitness and age-specific survival and fertility in a 19th century american population. Evolution.

Owen-Smith, N., Mason, D. R., Ogutu, J. O., 2005. Correlates of survival rates for 10 african ungulate populations: density, rainfall and predation. Journal of Animal Ecology 74 (4), 774–788.

Rice, S. H., 2004. Evolutionary Theory: Mathematical and Conceptual Foundations. Sinauer Associates Sunderland.

Roff, D. A., 1992. Evolution of Life Histories: Theory and Analysis. Springer.

Stearns, S., 1992. The Evolution of Life Histories. Oxford University Press.

Turelli, M., Petry, D., 1980. Density-dependent selection in a random environment: an evolutionary process that can maintain stable population dynamics. Proceedings of the National Academy of Sciences 77 (12), 7501–7505.

Voland, E., 1990. Differential reproductive success within the krummhörn population (germany, 18th and 19th centuries). Behavioral Ecology and Sociobiology 26 (1), 65–72.

